# Manipulation of *REMORIN* gene dosage affects salt signaling and tolerance in Arabidopsis thaliana

**DOI:** 10.64898/2025.12.22.695520

**Authors:** Michelle von Arx, Mariya Roussinova, Emmanuelle Bayer, Julien Gronnier

## Abstract

Salinity is among the most detrimental environmental plant stressors hampering growth, development and immunity. Its impact is increasing due to intensive agricultural practices and climate change. The molecular components linked to the early cellular and molecular events triggered by salt are mostly unknown. Here we show that genes encoding for members of the REMORIN proteins family exhibit dynamic temporal regulation following salt perception. Analogously to pattern-triggered immune signaling, we observed that salt rapidly induces the reorganization of REMORIN1.2 (REM1.2) into static plasma membrane nanodomains which colocalize with the actin-nucleating protein FORMIN 6. Overexpression of REM1.2 inhibits early signaling and late cell morphological responses to salt, defects which correlated with an increase in salt sensitivity. Our findings connect early salt-triggered signaling events to changes in cell architecture and root resilience and describe REMORIN nanodomains as a convergent articulation point in biotic and abiotic signaling pathways.

## Introduction

Soil salinity, predominantly caused by an excess of sodium (Na^+^) and chloride (Cl^-^) ions, severely limits agricultural productivity worldwide. Most of the major crop species are glycophytes (Zhao *et al*, 2021), and excess of salt in the soil affects their growth, development, reproduction, disease resistance and consequently impacts their overall fitness (Dinneny, 2010; Geng *et al*, 2013; Guo *et al*, 2022; Kang *et al*, 2008; Liu *et al*, 2015; Soltabayeva *et al*, 2021; van Zelm *et al*, 2020; Wang *et al*, 2021; Wu *et al*, 2021; Zhao *et al*., 2021). Over 20% of the cultivable areas worldwide are affected by soil salinity, a situation that is worsen at increasing pace by the global climate change and intensive agricultural practices, raising major concerns for food security globally (Arora, 2019; Mukhopadhyay *et al*, 2021). On the molecular scale, salt induces a two-fold stress. The accumulation of Na^+^ reduces water availability resulting in osmotic stress (Munns & Tester, 2008). Further, salt is toxic and exerts an ionic stress by disrupting cellular ionic homeostasis (Yang & Guo, 2018). Salt perception induces an array of early signaling events, such as the production of reactive oxygen species, the phosphorylation of MITOGEN-ACTIVATED PROTEIN KINASEs (MAPKs), and an influx of calcium (Ca^2+^), that culminate in a vast transcriptional reprogramming and changes in hormonal dosage (Geng *et al*., 2013; van Zelm *et al*., 2020). The best studied signaling cascade induced by the ionic stress is the salt overly sensitive (SOS) signaling pathway (Zhu, 2016) which is initiated by the sensing of the salt-triggered increase in cytoplasmic Ca^2+^ by the EF-hand calcium binding protein SOS3. The activated SOS3 binds to and activates the serine/threonine protein kinase SOS2 which in turn phosphorylates and activates the Na+/H+ antiporter SOS1, leading to Na^+^ detoxification (Ji *et al*, 2013; Ramakrishna *et al*, 2025). Much less is known regarding the identity of the molecular and cellular components engaged in early salt sensing and signaling events. The plasma membrane glycosyl inositol phosphoryl-ceramides (GIPCs) are proposed to bind Na^+^ and mediate the activation of salt-evoked Ca^2+^ permeable channel(s) (Jiang *et al*, 2019). The perception of the phytocytokine PAMP-INDUCED SECRETED PEPTIDE 3 (PIP3) by the RECEPTOR-LIKE PROTEIN KINASE 7 (RLK7) conditions salt-induced activation of the MAPK3 and MAPK6 (Zhou *et al*, 2022). Conversely, loss of the RAPID ALKALANIZATION FACTOR (RALF) peptides receptor *FERONIA* (*FER*) leads to enhanced salt-induced MAPK activation (Gigli-Bisceglia *et al*, 2022). Osmotic stress signaling has been shown to involve Rho of Plants GTPase-mediated activation of NADPH oxidases (Smokvarska *et al*, 2020), a process that is under control of FER-mediated regulation of phosphatidylserine (Smokvarska *et al*, 2023). Osmotic signaling also involves the plasma membrane localized C2 domain-containing protein BONZAI1 (Chen *et al*, 2020), and cytosolic RAF-SnRK2 kinase modules (Saruhashi *et al*, 2015; Takahashi *et al*, 2020). In the hours following salt application, salt-induced changes in the cell wall (Colin *et al*, 2022; Feng *et al*, 2018; Gigli-Bisceglia *et al*., 2022), and in the organization of the cytoskeleton (Kesten *et al*, 2019; Vilarrasa-Blasi *et al*, 2024; Wang *et al*, 2010), are linked with changes in cellular shape in the root of *Arabidopsis thaliana* (hereafter Arabidopsis) (Dinneny *et al*, 2008; Vilarrasa-Blasi *et al*., 2024). Changes in the cell wall are sensed by FER to mediate growth recovery (Feng *et al*., 2018), and wall-located RALF receptors, the LEUCINE RICH REPEAT EXTENSINs (LRXs), also condition salt tolerance (Zhao *et al*, 2020; Zhao *et al*, 2018). Whether early and late cellular and molecular responses to salt are linked is unknown.

REMORINs (REMs) are plant-specific structural components of the plasma membrane proposed to function as membrane scaffolds regulating the organization of plasma membrane lipids and proteins and influencing plasma membrane biophysics and topology (Gouguet *et al*, 2021; Gronnier *et al*, 2017; Huang *et al*, 2019; Legrand *et al*, 2023; Raffaele *et al*, 2009; Su *et al*, 2023). REMs have been divided into 6 groups (Raffaele *et al*, 2007) and are best characterized in the context of plant-microbe interactions and biotic stresses. In *Solanaceae* and in *Brassicaceae*, group 1 REMs regulate cell-cell movement of viruses (Fu *et al*, 2018; Huang *et al*., 2019; Jolivet *et al*, 2025; Raffaele *et al*., 2009). In Legumes, the symbiotic REM1 (SYMREM1) conditions the establishment of root nodulation (Lefebvre *et al*, 2010; Liang *et al*, 2018). In Arabidopsis, group 1 REMs mediate the nanodomain organization of the actin nucleator protein FORMIN 6 during pattern-triggered immunity signaling and contribute to the resistance to *Xanthomonas campestris* (Ma *et al*, 2021). In addition, Arabidopsis group 1 REMs have been linked to abscisic acid (Demir *et al*, 2013), salicylic acid (Huang *et al*., 2019), and auxin signaling pathways (Ke *et al*, 2021). Comparatively, the potential roles of REMs in abiotic stresses have been less studied. Interestingly, the heterologous overexpression of REM genes from salt-resistant poplar (*Populus euphratica*), from Foxtail Millet (*Setaria italica*) and from Mulberry (*Morus indica*) confer increase salt tolerance in Arabidopsis (Checker & Khurana, 2013; Yue *et al*, 2014; Zhang *et al*, 2020a), suggesting REMs play a role in salt stress tolerance.

Here we investigate the function of Arabidopsis REMs in salt stress. We show that *REMs* genes are dynamically regulated following salt perception and that their expression kinetics clusters in three groups. We observed that salt perception rapidly leads to the plasma membrane nanodomain organization of the group 1 REMs, REM1.2 and REM1.3, and that salt-induced REM1.2 nanodomains contains the actin nucleator protein FORMIN 6, indicating a role for REMs in the early signaling events associated with salt perception. We show that loss of the group 1 REMs *REM1.2, REM1.3* and *REM1.4* does not alter sensitivity to salt, while the overexpression of REM1.2 and REM1.3 is sufficient to inhibit salt-induced MAPK phosphorylation and tolerance to salt. Further we observed that changes in cellular morphology induced by salt are inhibited by the overexpression of REM1.2 and REM1.3. Our findings connect early salt-triggered signaling events to changes in cell architecture and root resilience and describe REMORIN nanodomains as a convergent articulation point in biotic and abiotic signaling pathways.

## Results

### Dynamic transcriptional regulation of *REMORINs* during salt stress

Plant responses to salt are highly dynamic processes, both temporally and spatially, as shown for instance by transcriptomic analyses (Dinneny *et al*., 2008; Wu *et al*., 2021). Analysis of microarray (Raffaele *et al*., 2007)(Supplementary Fig. S1A) and RNA sequencing experiments (Supplementary Fig. S1B) showed that several *REM* genes were responsive to salt treatment. The observed variability among experimental outcomes is likely attributable to differences in developmental stages, tissues, NaCl concentrations, and treatment durations used across studies. To investigate *REMs* transcripts accumulation in a time-resolved manner, we used temporal salt stress transcriptomic experiments (Wu *et al*., 2021). Interestingly, we observed that *REMs* showed dynamic transcript accumulation changes over time (Supplementary Fig. S2). For instance, the relative transcript accumulation of *REM4.1* was strongly upregulated 1h after salt application and subsequently downregulated a later timepoints (Supplementary Fig. S2). Conversely, *REM6.1*, *REM6.7* and *REM2.1* transcripts were downregulated at early timepoints but strongly upregulated 24h after salt treatment (Supplementary Fig. S2). We asked whether *REMs* showing distinct transcript accumulation profiles belong to separate functional clusters. To identify potential REM-associated functional clusters, we compared transcription accumulation profiles of each REM with the entire Arabidopsis transcriptome (Supplementary table 1). Based on K-means clustering, these analyses highlighted two to three main transcription clusters (Supplementary table 2), with distinct transcript accumulation profile dynamics (Fig. 1A and 1B). Cluster 1 is composed of *REM6.2*, *REM4.1*, *REM1.3* and *REM4.2* and 473 co-regulated genes which were transiently upregulated between 1 and 3 hours after salt treatment and downregulated at later timepoints. Similarly, the 241 co-regulated genes of cluster 2, which is composed of *REM3.2*, *REM3.1*, *REM6.6*, *REM5.1*, *REM6.5*, *REM1.2*, displayed a dynamic upregulation at early timepoints and a more drastic downregulation at later timepoints (Fig. 1B). Finally, the 151 co-regulated genes of Cluster 3, which includes *REM6.7*, *REM6.1*, *REM2.1*, were transiently downregulated between 1 and 3 hours and upregulated at later timepoints, thus presenting a profile opposite to cluster 1 and 2 (Fig. 1B). These transcriptional clusters showed distinct gene ontology (GO) enrichments (Supplementary Fig. S3), suggesting they may be linked to specific cellular and molecular functions. Together these analyses highlight the dynamics of *REMs* transcript accumulation following salt treatment and suggest REMs play a role salt stress response in Arabidopsis.

**Figure 1.**
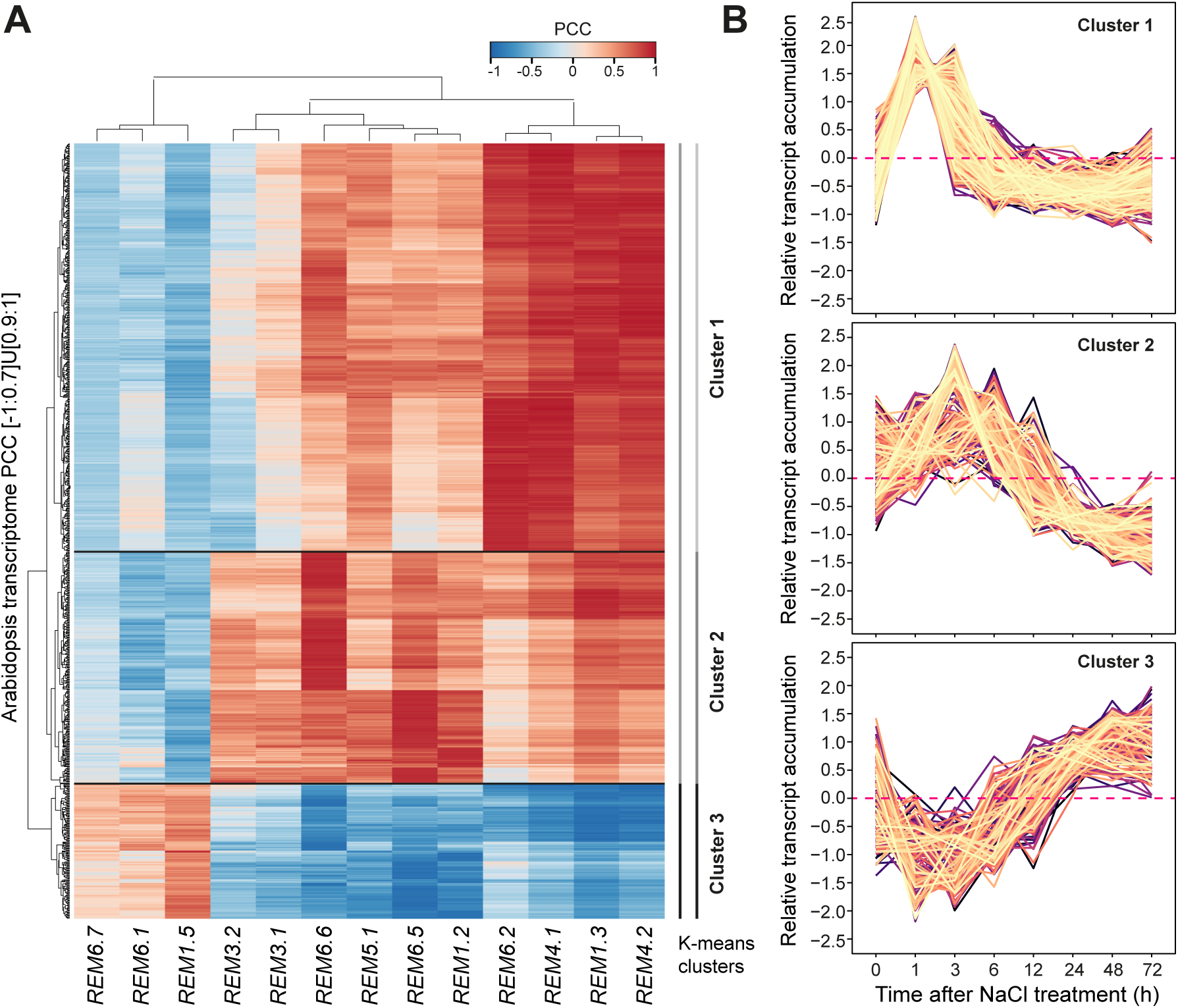
Dynamic transcriptional regulation of *REMORINs* during salt stress. **A**. Heat map of genes correlated with at least one REM (Pearson correlation coefficient ≥|0.9|). Temporal transcript accumulation profiles upon salt treatment cluster in two to three functional groups based on K-means clustering. **B**. Scaled transcript accumulation dynamics corresponding to each cluster.

### Salt induces plasma membrane nanodomain organization of REM1.2

The plasma membrane is dynamically organized into a myriad of nano-compartments, termed nanodomains (Jaillais *et al*, 2024). Heterologously expressed in *Nicotiana benthamiana,* REMs tend to form static nanodomains (Bücherl *et al*, 2017; Gronnier *et al*., 2017; Jarsch *et al*, 2014). In stable transgenic Arabidopsis lines REM1.2 appears dynamic and laterally diffuses within the plasma membrane (Jarsch *et al*., 2014; Jolivet *et al*., 2025). REM1.2 was shown to form stable nanodomains following the perception of the bacterial flagellin and was linked plant disease resistance (Ma *et al*., 2021). We ask whether salt perception would similarly lead to the reorganization of REM1.2. We used stable transgenic Arabidopsis lines expressing GFP and YFP-tagged REM1.2 under its native promoter in a *rem1.2* knock-out background (Huang et al., 2019; Jarsch et al., 2014) (Supplementary Fig. S4). Using laser scanning confocal microscopy, we examined the plasma membrane organization of REM1.2 in root epidermal cells. In good agreement with previous studies (Jarsch *et al*., 2014; Jolivet *et al*., 2025; Ma *et al*., 2021), we found that under control conditions REM1.2 showed a diffuse pattern at the PM (Fig. 2A). In contrast, within 2 minutes of 140 mM NaCl treatment we observed a re-organization of YFP-REM1.2, YFP-REM1.3 and GFP-REM1.2 into static and stable nanodomains (Fig. 2, Supplementary Fig. S5-7). Indeed, we observed that 140 mM NaCl led to the appearance of YFP-REM1.2 nanodomain to an average density of 0.6 nanodomain per square microns (Fig. 2C) and to an increase in the image-wide spatial clustering index (Fig. 2D), which were not linked to global changes in fluorescence intensity at the plasma membrane (Fig. 2E), suggesting that YFP-REM1.2 nanodomains emerges from lateral re-organization.

**Figure 2.**
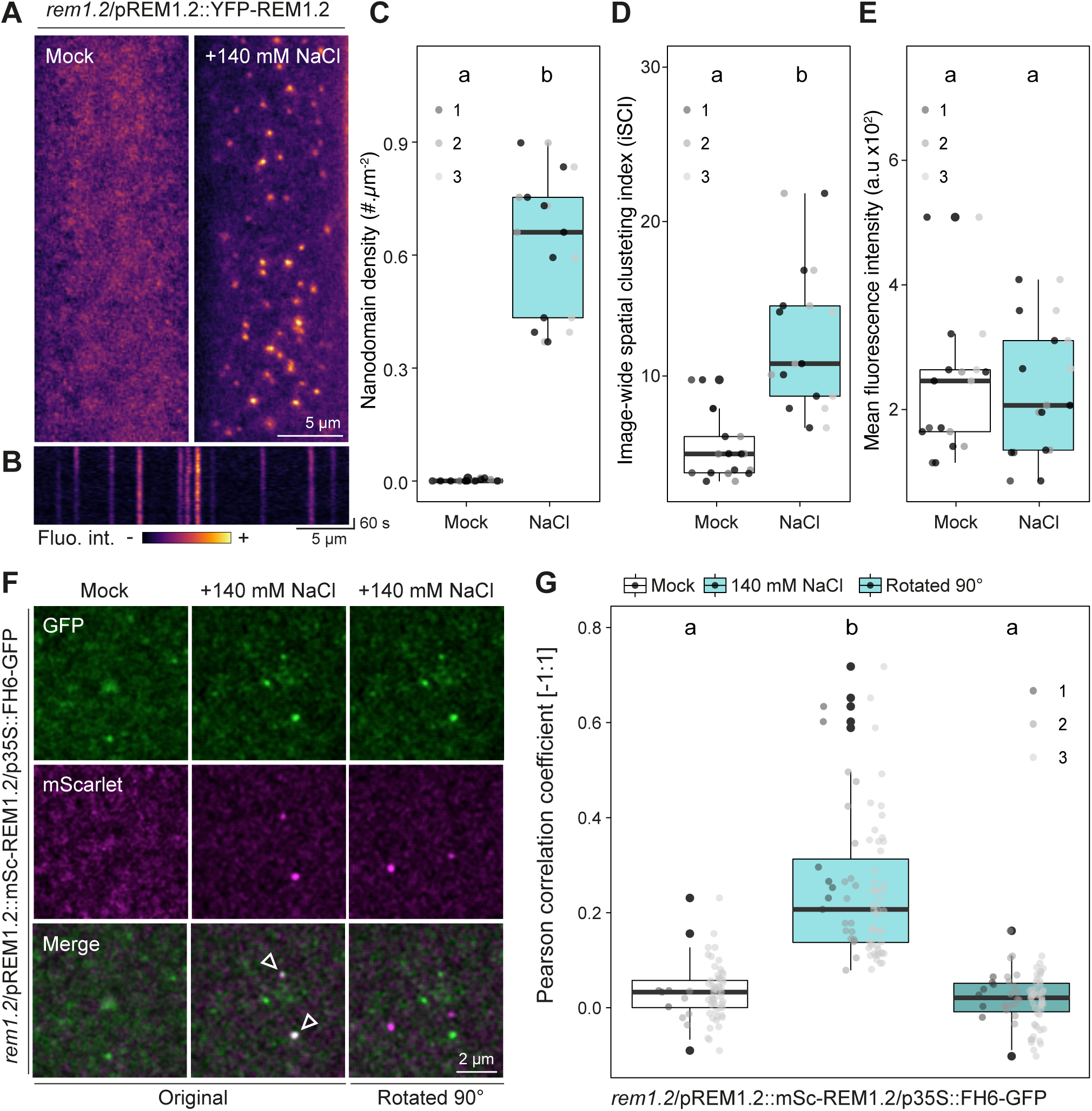
Salt triggers the re-organization of REM1.2 into plasma membrane nanodomains containing the actin nucleator protein FORMIN 6. **A**. Confocal microscopy pictures of the cell surface of root epidermal cells in presence and in absence of NaCl 140 mM, observed within two minutes of treatment. **B**. Temporal projection of YFP-REM1.2 nanodomains upon treatment with 140mM NaCl. **C-E**. Quantification of YFP-REM1.2 nanodomain density per µm^2^ (**C**), of YFP-REM1.2 image wide spatial clustering index (iSCI) (**D**), and of YFP-REM1.2 mean image fluorescence intensity (**E**) in presence of 140 mM NaCl treatment or corresponding mock control. Each dot represents the mean measurement per plant, dot colors indicate independent experiments, total number of seedlings and cells (n,n) for mock (n=9, n=82) and (n=9, n=81) for NaCl. Conditions which do not share a letter are significantly different in pairwise Wilcoxon test with Bonferroni p-value adjustment (p< 0,05). **F**. Confocal microscopy images of mScarlet-REM1.2 and FH6-GFP in 5 days old seedlings cotyledon epidermal cells observed between one and ten minutes of 140 mM NaCl treatment or corresponding mock control. **G**. Pearson correlation coefficient analysis of the co-localization between mScarlet-REM1.2, FH6-GFP in presence of 140 mM NaCl treatment or corresponding mock control. 90°C rotated images of the mScarlet channel (**F**) were used as randomized images for comparative analysis. Each dot represents the correlation coefficient obtained from one individual Cell and colors indicate individual experiments. Conditions which do not share a letter are significantly different in pairwise Wilcoxon test with Bonferroni p-value adjustment (p< 0,05). Total number of Cells (n): mock (47), NaCl (31), control (31).

We next wondered whether salt-induced nanodomain re-organization was specific to root epidermal cells. We observed that NaCl treatment led to the organization of REM1.2 in cotyledon and hypocotyl epidermal cells (Supplementary Fig. S6 and S7), suggesting it corresponds to a conserved cellular response of the epidermis. In hypocotyl, YFP-REM1.2 nanodomains of distinctive sizes appeared over time (Supplementary Fig. S6) which seemed to emanate from the lateral sliding and assembly of smaller nanodomains (Supplementary Fig. S6, Supplementary movie 1) and were not observed in root or cotyledon epidermal cells. We next questioned whether the osmotic stress component of salt would be sufficient to promote the nanodomain organization of REM1.2. To compare with the nanodomain organization observed upon 140 mM NaCl, we used iso-osmotic concentration of sorbitol (280 mM), a non-electrolyte which thus only induces a comparable osmotic stress. We observed that as for NaCl, sorbitol treatment led to the reorganization of GFP-REM1.2 into nanodomains in both root and cotyledons epidermal cells (Supplementary Fig. S7). This indicates that the osmotic component of NaCl is sufficient to promote REM1.2 nanodomain organization. Interestingly, in control conditions REM1.2 appeared organized in nanodomains in a subset of guard cells (Supplementary Fig. S8). These observations may be linked to stomata water potential and potentially linked to the regulation of REM1.2 nanodomain-hosted ABA signaling and of stomatal closure (Demir *et al*., 2013).

REM1.2 and REM1.3 were shown to be re-organized into plasma membrane nanodomains during pattern-triggered immune signaling and thereby condition the nanodomain organization and the activity of the actin nucleating protein FORMIN 6 (FH6) (Ma *et al*., 2021). We therefore wondered whether salt-induced nanodomain organization of REM1.2 would also be linked to FH6. We use a stable transgenic line co-expressing mScarlet-REM1.2 and FH6-GFP (Ma *et al*., 2021) to analyze their potential co-localization. In accordance with a previous report (Ma *et al*., 2021), in the absence of any stimuli, no particular co-localization was observed between FH6-GFP and mScarlet-REM1.2 (Fig. 2F and 2G). In contrast, minutes after the application of 140 mM NaCl we observed that a subpopulation of salt-induced mScarlet-REM1.2 nanodomains co-localized with FH6-GFP nanodomains (Fig. 2F and 2G). These observations suggest that, similar to its role in pattern-triggered immune signaling, REM1.2 may regulate FORMIN plasma membrane organization and activity during salt signaling.

### Mis-regulation of *REM1.2* expression inhibits salt signaling and tolerance

The transcriptomic analyses (Fig. 1) and live cell imaging experiments (Fig. 2) suggest that REMs exert a role during salt perception. REM1.2 and REM1.3 are genetically redundant and loss of both *REM1.2* and *REM1.3*, but not of the individual genes, leads defect in passive cell-cell diffusion in Arabidopsis roots (Huang *et al*., 2019), in disease resistance against *Xanthomonas campestris pv. campestris* (Ma *et al*., 2021) and in the cell-cell movement of the Plantago asiatica mosaic virus (PlAMV) (Jolivet *et al*., 2025). Introducing a loss of function allele for *REM1.4* in *rem1.2/rem1.3* further aggravated PlAMV infection (Jolivet *et al*., 2025). We therefore used this characterized *REMs* triple mutant (*rem1.2/rem1.3/rem1.4*) for functional assays. As previously reported (Huang *et al*., 2019), the T-DNA mutant for *REM1.3* (salk_117448.53.95.x) presents residual accumulation of REM1.3 when combined with loss of *REM1.2* (salk_117637.50.50.x) (Supplementary Fig. S4). We assessed root growth of WT and of *rem1.2/rem1.3/rem1.4* mutant seedlings transferred to ½ MS supplemented with NaCl or corresponding mock control. As a positive control, we used a loss-of-function mutant allele for *FERONIA*, *fer-4* (Duan *et al*, 2010). As previously reported (Feng *et al*., 2018), we observed that *fer-4* root growth is greatly reduced compared to WT in medium containing 100 mM NaCl (Supplementary Fig. S9) and 140 mM NaCl (Supplementary Fig. S9). In contrast, the loss of *REM1.2*, *REM1.3* and *REM1.4* did not affect root sensitivity to 100 mM NaCl (Supplementary Fig. S9) and 140 mM NaCl (Supplementary Fig. S9). To corroborate these observations, we tested salt-induced phosphorylation of the MAPKs (Ichimura *et al*, 2000; Nakagami *et al*, 2005; Teige *et al*, 2004; Yu *et al*, 2010). We observed that loss of *REM1.2*, *REM1.3* and *REM1.4* did not affect salt-induced MAPK phosphorylation (Supplementary Fig. S9). These observations indicate either that despite being dynamically transcriptionally regulated, these REMs do not contribute to early salt signaling and root growth tolerance to salt, or that loss of *REM1.2*, *REM1.3* and *REM1.4* may be compensated by additional REMs. The transcriptional signatures of REM1.2 and REM1.3 are shared with at 8 additional REMs that belong to distinct groups (Fig. 1) and suggest that a high degree of genetic complexity safekeeps REM-associated function(s) in salt stress.

*REM1.2* and *REM1.3* are transiently upregulated before being downregulated (Fig. 1, Supplementary Fig. S2). We reasoned that miss-regulation of this pattern may affect salt tolerance. We use a transgenic line that conditionally overexpressed REM1.2 tagged with the fluorescent protein mRuby using the β-estradiol–inducible system (*XVE::mRuby-REM1.2*) (Ma *et al*., 2021). We assessed the root growth of *XVE::mRuby-REM1.2* in presence and absence of NaCl and β-estradiol, and of their corresponding mock controls. We observed that the overexpression of mRuby-REM1.2 led to a decrease in root growth upon salt treatment (Fig. 3A), indicating that REM1.2 overexpression increases salt sensitivity. Likewise, we observed that REM1.2 overexpression led to a reduction in root growth upon sorbitol treatment (Supplementary Fig. S7). Further, we observed that the overexpression of mRuby-REM1.2 inhibits salt-induced phosphorylation of the MAPKs (Fig. 3B). To corroborate these observations, we tested the effect of the conditional overexpression of untagged REM1.2 and REM1.3 driven by the β-estradiol–inducible system (Huang *et al*., 2019). We observed that the overexpression of REM1.2 and of REM1.3 inhibited salt-induced MAPK phosphorylation (Supplementary Fig. S10).

**Figure 3.**
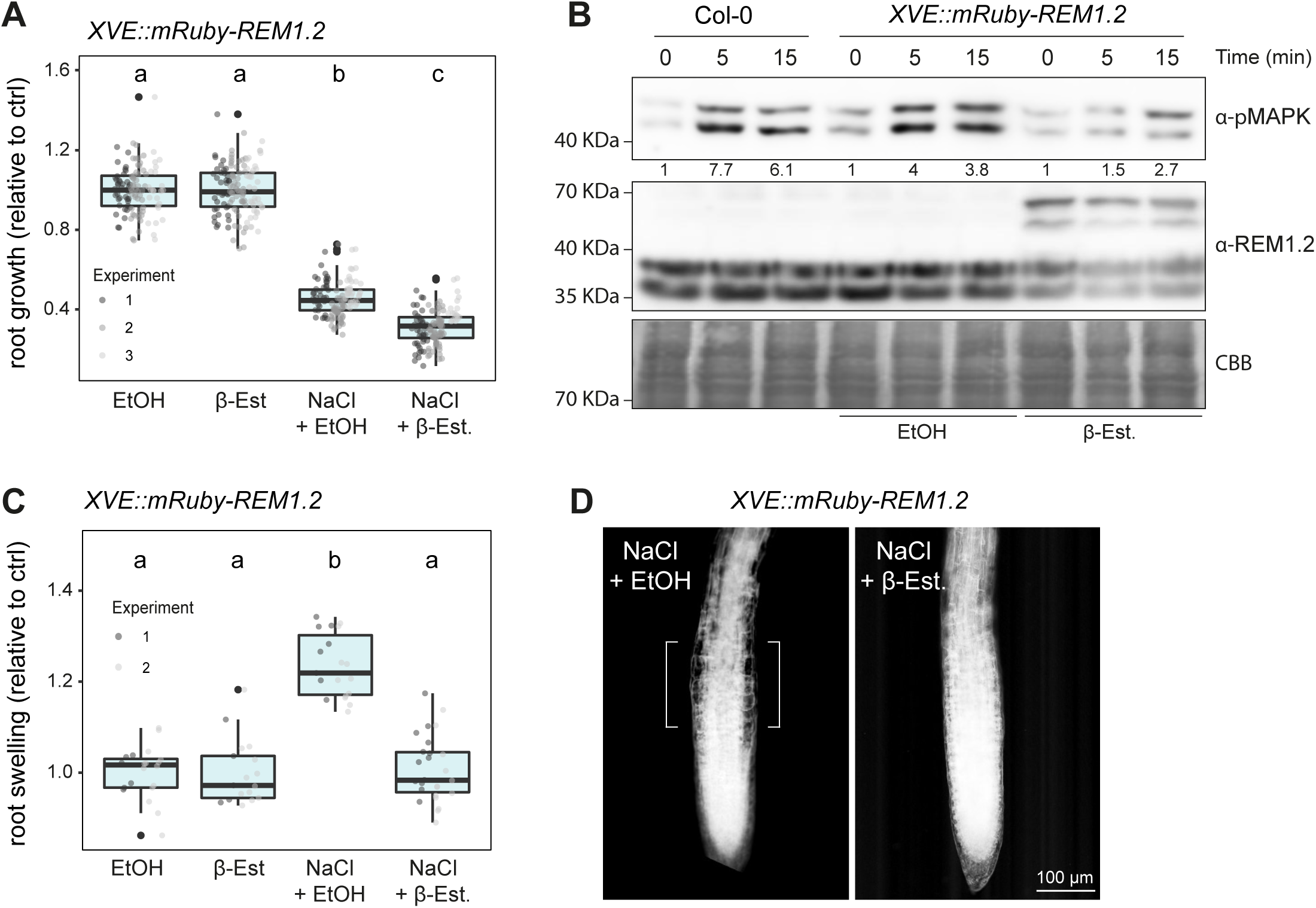
REM1.2 over-expression inhibits salt signaling and tolerance. **A**. Relative root growth measurements of seedlings transferred to ½ MS containing 0.5 μM β-Estradiol and/or 140 mM NaCl and/or corresponding controls for five days. Each dot represents measurements from individual seedlings, colors indicate individual experiments. Measurements are normalized to corresponding mock conditions (without NaCl). Conditions which do not share a letter are significantly different in pairwise Wilcoxon test with Bonferroni p-value adjustment (p< 0,05). Between 110 and 128 roots were measured in total. **B**. Western blot analysis of salt-induced MAPK phosphorylation in 10 days old seedlings treated with 140 mM NaCl in time course experiments. Western blots were probed with α-p44/42-ERK revealing phosphorylated MAPKs or α-REM1.2. Blot stained with Coomassie brilliant blue (CBB), is presented to show equivalent protein loading. Similar results were obtained in two independent experiments. **C**. Quantification of the root swelling of seedlings incubated in liquid ½ MS containing 0.5 μM β-Estradiol and/or 140 mM NaCl and/or corresponding controls for 24h. Each dot represents one measurement from one root, colors indicate independent experiments. Measurements are normalized to corresponding mock conditions (without NaCl). Conditions which do not share a letter are significantly different in pairwise Wilcoxon test with Bonferroni p-value adjustment (p< 0,05). Between 17 and 43 seedlings were analyzed in total. **D**. Representative confocal light transmission images (inverted grey values) of the root tip of 5 days old seedlings incubated in media 140 mM NaCl in absence and presence of 0.5 μM β-Estradiol.

The ionic and osmotic stresses induced by salt affects plant cell morphology on different but interconnected levels. Osmotic stress has been reported to induce changes in cytoskeletal organization, growth arrest and altered root-tip morphology (Geng *et al*., 2013; Kang *et al*., 2008; Vilarrasa-Blasi *et al*, 2021; Zhao *et al*., 2021). The ionic component of salt stress has been proposed to directly compromise cell wall integrity by disrupting pectin cross-linking (Feng *et al*., 2018; Wang *et al*., 2021). Upon pattern-triggered immune signaling, the overexpression of REM1.2 is sufficient to promote actin nucleation activity (Ma *et al*., 2021). We observed that salt triggered the formation of REM1.2 nanodomains that contain FH6 (Fig. 2), presumably promoting actin nucleation. The tight regulation of cytoskeletal organization is an important determinant to mitigate salt-induced cell morphological changes and subsequently root growth recovery (Feng *et al*., 2018; Geng *et al*., 2013; Vilarrasa-Blasi *et al*., 2021). We inquired whether the changes in cell morphology induced by salt could be altered by the overexpression of REM1.2. As previously observed in WT seedlings (Dinneny *et al*., 2008; Vilarrasa-Blasi *et al*., 2024), in absence of REM1.2 overexpression we observed that salt induces the swelling of the epidermal cells of the root transition zone (Fig. 3C-D, Supplementary Fig. S11). Interestingly, we observed that the overexpression of mRuby-REM1.2 inhibited salt-induced cell swelling (Fig. 3C-D). Likewise, we observed that the overexpression of untagged REM1.2 and REM1.3 inhibited salt-induced cell swelling (Supplementary Fig. S11). Altogether, these observations indicate that miss-regulation of REM1.2 and of REM1.3 gene dosages affect early and late responses to salt and indicate that a dynamic *REMORIN* gene dosage underlies salt signaling and tolerance in Arabidopsis thaliana.

## Discussion

Salt directly impacts plant growth and development leading to important crop yield losses. While some plant adaptative strategies have been described, much remains to be understood as to how plants sense salt, how plants coordinate salt stress responses, and on the identity of the cellular and molecular components underlying these processes. Here we show that REMs are transcriptionally dynamically regulated following salt sensing (Fig. 1), and that REM1.2 and REM1.3 rapidly form salt-induced nanodomains (Fig. 2, Supplementary Fig. S5-7) suggesting an active role for these proteins in early salt signaling. We observed that the overexpression of REM1.2 and REM1.3 inhibits salt-induced MAPK phosphorylation (Fig. 3) suggesting that they function as negative regulators of salt-induced signaling. We observed that the loss of *REM1.2*, *REM1.3* and *REM1.4* is not sufficient to alter root growth sensitivity to salt or salt-induced signaling (Supplementary Fig. S9) suggesting that additional REMs may compensate for their loss. This putative functional compensation may involve additional group 1 REMs (REM1.1 and REM1.5), the REM members we found co-regulated with REM1.2 and REM1.3 (8 members) and their close paralogues. Further research will be required to genetically dissect the potential role(s) of REMs in salt signaling and salt tolerance. While our data clearly show that the overexpression of REM1.2 inhibits root growth tolerance to salt and osmotic stress, the overexpression of REMs homologues from salt-resistant plant species promotes salt tolerance in Arabidopsis (Checker & Khurana, 2013; Yue *et al*., 2014; Zhang *et al*., 2020a). It will be interesting to define the potential molecular bases for the opposite effect of REM overexpression on salt tolerance. It is interesting to note that our analyses highlight anti-coregulated transcriptional clusters (Fig. 1) which may indicate that individual Arabidopsis REMs have antagonistic roles during salt stress. Defects in salt-induced MAPK phosphorylation have been linked with increase in salt sensitivity (Yu *et al.,* 2010; Zhou *et al,* 2021; Gigli-Bisceglia *et al*., 2022). It will be interesting to define whether the inhibition of salt-induced MAPK phosphorylation by REM1.2 and REM1.3 is linked to known regulatory components of MAPKs, such as Phospholipase D-mediated production of phosphatidic acid (Yu *et al*., 2010), the PIP3-RLK7 module (Zhou *et al*, 2021), FER-mediated cell wall signaling (Gigli-Bisceglia *et al*., 2022), or to unknown regulator(s). Further research will also be required to define whether REM1.2 and FH6-containing nanodomains (Fig. 2) regulate actin dynamics and whether this is linked to the effect of REM1.2 on early salt-triggered signaling and salt-induced cell morphological changes (Fig. 3) and whether parts or the combination of these observations explain the effect of REM1.2 on root growth tolerance to salt. Since both flg22-induced and salt-induced REM1.2 nanodomains contain the actin nucleating protein FORMIN6, and that REM1.2 has been shown to promote actin nucleation activity of FORMIN6 (Ma *et al.,* Plant Cell 2021), it is tempting to speculate that REM1.2 overexpression densifies the actin cytoskeleton, thereby inhibiting salt-induced cell swelling. In line with our observations, a recent pre-print reports that an hyperosmotic stress triggers the re-organization of REM1.2 into nanodomain and identified additional molecular actors associated with REM1.2 during this process (Rui *et al*, 2025), pointing toward the formation of osmotic stress-induced supra-molecular complexes.

Our observations notably link the absence of salt-induced cell swelling with an increase salt sensitivity (Fig. 3, Supplementary Fig.S11). While the defect in salt-induced MAPK phosphorylation could be sufficient to explain the effect of REM1.2 overexpression, it may also be interesting to consider that cell swelling and the sensing of the associated modifications of cell mechanics may be required to mount a full adaptative response. Interestingly, root swelling was proposed to correspond to an adaptative mechanism conditioning stress tolerance in Maize (Byrt *et al*, 2018; Li *et al*, 2014). In addition, it will be interesting to define whether the regulation of salt sensitivity by REMs is linked to the regulation of H^+^-ATPases (Xhelilaj et al., 2025). At the cellular level, salt notably induces, cell plasmolysis, a bulk endocytosis-meditated internalization of plasma membrane material (Baral *et al*, 2015; Ueda *et al*, 2016), the depolymerization of actin and microtubule cytoskeleton (Kesten *et al*., 2019; Lian *et al*, 2025; Wang *et al*., 2010) and the re-localization of the CELLULOSE SYNTHASE 6 and COMPANION OF CELLULOSE SYNTHASE1 to post-Golgi-related compartments (Endler *et al*, 2015). These events have been described to occur from 15-30 minutes following salt treatment, while salt-induced changes in cell morphology are observed 4-6 hours after salt treatment (Feng *et al*., 2018). We observed that salt induces REM1.2 organization within 2 minutes following treatment. Further, we observed that an osmotic stress was sufficient to trigger REM1.2 nanodomain organization. Similarly, osmotic stress induces the nanodomain organization of ROP6 (Smokvarska *et al*., 2020) and the receptor-like kinase QIAN SHOU KINASE (QSK1) (Grison *et al*, 2019) within minutes. Altogether, these observations indicate that a rapid and comprehensive plasma membrane re-organization is triggered by the osmotic component of salt stress. To our knowledge, these events correspond to the earliest cell structural modifications triggered by salt and may relate to the formation of several co-existing structural and functional units involved in salt sensing and signaling.

## Acknowledgements

We thank all present and past members of the NanoSignaling Lab for fruitful discussion and comment on the manuscript. We thank Thomas Ott, Yansong Miao and Xu Chen for sharing published material. Confocal imaging was performed at the ZMBP microscopy facility and at the Center for Advanced Light Microscopy (CALM) of the TUM school of Life Sciences. This research was supported by the Deutsche Forschungsgemeinschaft (DFG) grants (A08-SFB1101 and B01-TRR356) to JG.

---

**Supplementary table 1 | Gene expression correlation**

**Supplementary table 2 | REMORIN-associated transcriptional clusters**

**Supplementary movie 1 | Lateral diffusion of YFP-REM1.2 nanodomains in hypocotyl epidermis treated with NaCl 140 mM**

## Material and methods

### Plant material and growth

*Arabidopsis thaliana* ecotype Columbia (Col-0) was used as WT control. The *fer-4* (Duan *et al*., 2010), *rem1.2/rem1.3/rem1.3* (Jolivet et al., 2025), *rem1.2*/*pREM1.2::YFP-REM1.2*, *rem1.3*/*pREM1.3::YFP-REM1.3* (Jarsch *et al.,* 2014), *rem1.2*/*pREM1.2::GFP-REM1.2, XVE::REM1.2*, *XVE::REM1.3* (Huang *et al*., 2019), *XVE::mRuby-REM1.2* and *rem1.2*/*pREM1.2::mScarlet-REM1.2*/*p35S::FH6-GFP* (Ma *et al*., 2021) lines were previously reported. Seeds were surface sterilized using chlorine gas for 5 h or by incubating them in 0.1 % Tween20 in 70 % EtOH for 10 min, following 70 % EtOH for 10 min and 100 % EtOH for 1min. Seeds were stratified for 2 days in the dark at 4 °C and grown on half Murashige and Skoog (MS) media pH 5.8 supplemented with vitamins, 1 % sucrose and 0.8 % agar (Duchefa Biochemie, 9002-18-0), at 22 °C and a 16-h light photoperiod.

### Analysis of root growth

For root growth measurements surface-sterilized seeds were placed in a straight line and grown vertically on square petri-dishes. All seeds were stratified for 2 days in the dark at 4 °C and then placed in a controlled growth chamber (22 °C, 16 h light) for 3 days. Seedlings from one plate were divided and transferred to fresh media containing treatment or respective controls, the position of their root tip was marked on the plate. 4 days after transfer, the seedlings were imaged using a conventional camera, and the relative root growth from marked position to the root tip were measured using the Fiji (Schindelin et al. 2012). Salt experiments were conducted using 100 mM or 140 mM NaCl final concentration from a stock solution of 5M NaCl. Sorbitol experiments were conducted using 280 mM sorbitol final concentration. For estradiol induction, a concentration of 0.5 µM or 5 µM β-Estradiol in EtOH or corresponding volume of 100 % EtOH as control we used.

### MAPK phosphorylation assays

For MAPK phosphorylation assays, 7-day old seedlings were grown vertically on solid ½ MS plates and transferred to grow for one week in 6 well plates containing liquid ½ MS supplemented with either 0.5 or 5 μM β-Estradiol or corresponding volume of 100 % EtOH as control. For one sample, four seedlings from 2 individual wells were treated with 140 mM NaCl for the indicated time and flash frozen in 2 mL tubes containing glass beads in liquid nitrogen.

### Protein extraction and immunoblotting

Protein extraction and immunoblotting were performed as previously described (Biermann *et al*, 2025; Gronnier *et al*, 2022). Samples were ground for 30 seconds at a frequency of 25 movement per second to a fine powder using TissueLyser II (QIAGEN). 200 μL protein extraction buffer was added, and samples were incubated on a rotator for 40 min at 4 °C followed by a centrifugation for 10 min at 20 000 g at 4°C. The supernatant was transferred to new 1.5 mL tubes containing 40 μL of 6X SDS loading dye supplemented with 2 μL DTT and heated for 10 min at 90 °C. 15 μL sample were loaded on a 10 % SDS-PAGE gel and run at 150 volts for 60 min. Samples were transferred to activated PVDF membranes by wet transfer in Tris-Glycine buffer at 100 volts for 60-90 min at 4 °C. The membranes were blocked for one hour at room temperature with a 5 % milk-TBS 0.01 % Tween 20 solution and incubated in primary antibody solution. Polyclonal anti-REM1.2 (reference number C12-77) and polyclonal anti-REM1.3 (reference number C12-78) antibodies were raised in rabbit by CovalAb (France) against full length 6xHis-REM1.2 and 6xHis-REM1.3 proteins purified from E. coli. The rabbit anti-REM1.2 and the rabbit anti-REM1.3 used at 1:3000 5 % milk-TBS 0.01 % Tween 20 solution for 2 h at room temperature. The phospho-p44/42 MAPK (Erk1/2) (Thr202/Tyr204) antibody (Cell Signalling Technology, #9101) was used at 1/4000 in 5 % BSA TBS 0.01 % Tween 20 solution for 2 h at room temperature. Following three 10-min washes in 5% milk-TBS 0.01 % Tween 20, the secondary peroxidase-couple anti-rabbit (Sigma, A0545-1ML) was used at 1:10 000 5% milk-TBS 0.01 % Tween 20 solution for 1 h. After washing two times in TBS 0.01% Tween 20, the membranes were revealed using SuperSignal™ West Pico PLUS Chemiluminescence Substrate (Thermo Scientific) and a ChemiScope 6000 Series (CLINX). To assess protein loading, membranes were stained in Coomassie blue brilliant (CBB) staining solution.

### Confocal laser scanning microscopy and image analyses

Seedlings used for microscopy experiments were surface sterilized, sowed in a straight line on agar ½ MS plates, stratified for 2 days in the dark at 4 °C, then placed in a controlled growth chamber at 22°C and kept under a 16-hour photoperiod under environmentally controlled for 5 days. Confocal microscopy was performed as previously described (Gronnier *et al*., 2022). The experiments were conducted using a Leica SP8 CLSM system (Leica, Wetzlar, Germany) equipped with Argon, DPSS, He-Ne lasers and hybrid detectors, or a Zeiss LSM 880 equipped with Argon, DPSS lazers and GaAsP detectors. The following excitation wavelength 488 nm (GFP), 514 nm (YFP) and 561 nm (mScarlet) and fluorescence emission wavelength 495-550 nm (GFP), 520-550 nm (YFP) and 571-651 nm (mScarlet) were used. To analyse the plasma membrane organization of REM1.2 and REM1.3 localized proteins, 5-day-old seedlings were mounted in ½ MS liquid media in absence or presence of indicated treatment or corresponding control solution, between glass slide and coverslip. The experiments were performed using strictly identical confocal acquisition parameters (e.g. laser power, gain, zoom factor, resolution, and emission wavelengths reception), with detector settings optimized for low background and no pixel saturation. Pseudo-color images were obtained using look-up-table (LUT) in Fiji (Schindelin *et al*, 2012). The quantification of REM1.2 and of REM1.3 plasma membrane organization was performed using NanoNet accessible at Github (https://github.com/NanoSignalingLab/NanoNet). Arabidopsis transgenic line *rem1.2*/*pREM1.2::mScarlet-REM1.2*/*p35S::FH6-GFP* (Ma *et al*., 2021) was imaged using a 63X objective (HC PL APO CS2) and excitation wavelength 488 nm (GFP) and 561 nm (mScarlet) and fluorescence emission was collected between 495-550 nm (GFP) and 571-651 nm (mScarlet). Pearson correlation coefficients were determined using the JaCop plugin (BOLTE & CORDELIÈRES, 2006) in Fiji (Schindelin *et al*., 2012).

### Analysis of salt-induced root swelling

5 days old seedlings were transferred in liquid media contain 140 mM NaCl or corresponding mock solution (1/2 MS), 13h prior imaging. The roots were imaged using a 10X dry objective (HC PL APO CS2) and obtained images in brightfield mode. Relative swelling was quantified by diving the width of transition zone (salt-induced swelling zone) by the width the root 2 mm above this zone.

### Cloning, heterologous protein expression and purification

The coding sequences of REM1.2 and REM1.3 were amplified from *Arabidopsis thaliana* Col-0 cDNA and cloned into the pDONR211 and subcloned into the pDEST17 for 6xHis fusions by Gateway recombination (Thermo Scientific). 6xHis-REM1.2 and 6xHis-REM1.3 recombinant proteins were expressed in *Escherichia coli* BL21 DE3 cells and purified using fast flow chelating sepharose resin (Amersham) according to manufacturer’s instructions as previously described (Perraki *et al*, 2018).

### Analysis of transcriptomic data

The procedure and R code R generated and used to extract and normalize reads as well as to compute the gene correlation matrix and transcriptomic profile over time can be found on Github (https://github.com/NanoSignalingLab/NanoNet). Briefly, raw read counts of the experiment “GSE153103” were retrieved from the NCBI GEO repository (<u>GSE153103</u>, https://www.ncbi.nlm.nih.gov/geo/). Gene lengths were obtained from Arabidopsis reference genome (https://www.arabidopsis.org/) and used to normalize the transcriptomic data. Differential gene expression was calculated using the DESeq2 package (Love *et al*, 2014) from Bioconductor (https://bioconductor.org/) in R (https://www.R-project.org/). Genes that were correlated by Pearson correlation coefficient >|0.9| with at least one REM were then filtered. The transcript accumulation correlation matrix was then clustered and re-organized using the heatmap.2 function from the gplots package in R with default Euclidian distance. The normalized Fkpm temporal transcriptomic profile of gene clusters together with the respective REMs were plotted using R. Gene Ontology enrichment analyses were performed using ShinyGO (http://bioinformatics.sdstate.edu/go/). To visualize gene expression profiles over time, transcript accumulation for each gene was normalized by subtracting the mean and dividing by the standard deviation using the default base::scale() function in R. For micro array data, the three datasets GSE264404 (Cho, 2024), GSE32087 (Nishiyama *et al*., 2012), GSE94885 (Abdelrahman *et al*., 2021), were retrieved from the NCBI GEO repository, filtered for REMs and plotted in R. For Meta analysis of different RNA sequencing experiments, the following datasets were retrieved from the Gene Expression Omnibus and the European Nucleotide archive. PRJEB33124 (Kawa *et al*., 2020), PRJNA290554 (Van Tol *et al*., 2019), PRJNA295091 (Suzuki *et al*., 2016), PRJNA344695 (Carrasco-López *et al*., 2017), PRJNA439011 (Palomar *et al*., 2021), PRJNA476945 (Shkolnik *et al*., 2019), PRJNA478206, PRJNA513852 (Blanco-Touriñán *et al*., 2021), PRJNA525176 (He *et al*., 2019), PRJNA544786 (Zhang *et al*., 2020b), PRJNA544879, PRJNA554020 (Keshishian *et al*., 2022), SRP000935 (Filichkin *et al*., 2010). Datasets were filtered for REMs, scaled for each experiment and plotted using R.

## Data analysis and statistics

The number of independent experiments and the number of individual cells or root analyzed per condition and collected across these experiments are indicated in each figure legends. All statistical test and data visualization were performed in R (https://www.R-project.org/) using the statistical test indicated in the figure legends.

## Code availability

All Fiji macros, R scripts and python scripts were made publicly available at the following Github (https://github.com/NanoSignalingLab).

---

**Supplementary Figure S1.**
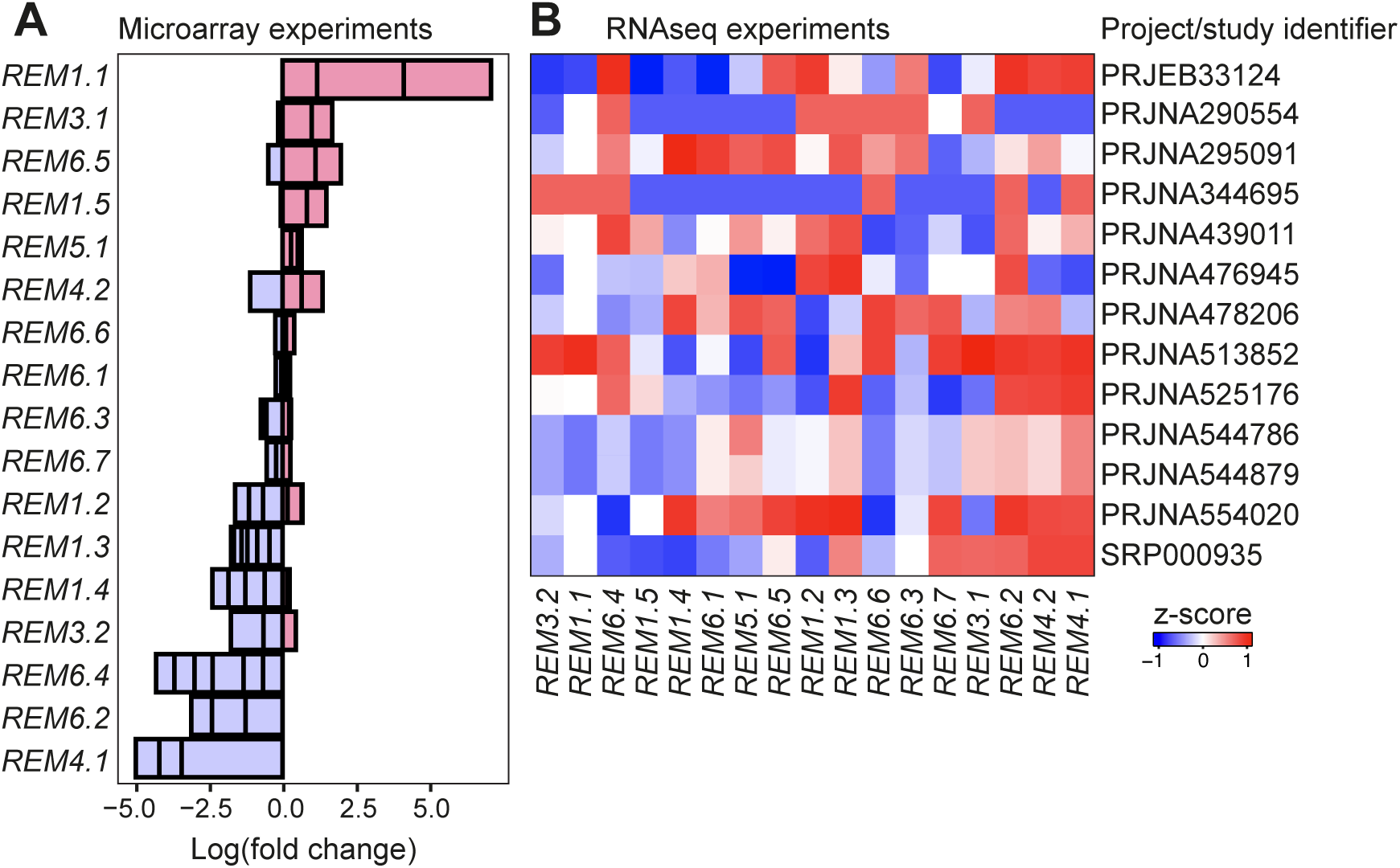
Analysis of *REMs* transcript accumulation in microarrays and RNA sequencing experiments. **A**. Fold change REMs transcript accumulation upon NaCl treatment in the microarray experiments GSE264404 (Cho, 2024), GSE32087 (Nishiyama *et al*, 2012), GSE94885 (Abdelrahman *et al*, 2021), **B**. *REMs* expression z-score upon NaCl treatment in the RNA sequencing experiments PRJEB33124 (Kawa *et al*, 2020), PRJNA290554 (Van Tol *et al*, 2019), PRJNA295091 (Suzuki *et al*, 2016), PRJNA344695 (Carrasco-López *et al*, 2017), PRJNA439011 (Palomar *et al*, 2021), PRJNA476945 (Shkolnik *et al*, 2019), PRJNA478206, PRJNA513852 (Blanco-Touriñán *et al*, 2021), PRJNA525176 (He *et al*, 2019), PRJNA544786 (Zhang *et al*, 2020b), PRJNA544879, PRJNA554020 (Keshishian *et al*, 2022), SRP000935 (Filichkin *et al*, 2010). The data were retrieved from the Gene Expression Omnibus and the European Nucleotide Archive.

**Supplementary Figure S2.**
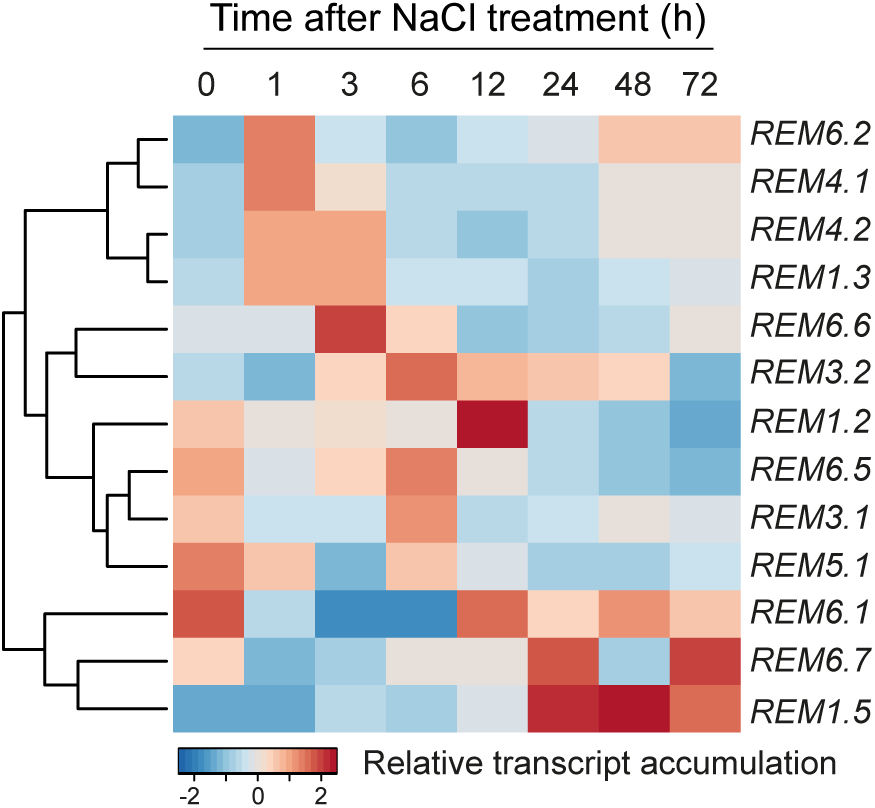
Analysis of *REMs* transcript accumulation in salt treatment time course experiment. Heatmap of normalized transcript accumulation dynamics of REMs upon salt stress over time. Each timepoint corresponds to the mean of 3 biological replicates from publicly accessible data by (Wu et al. 2021).

**Supplementary Figure S3.**
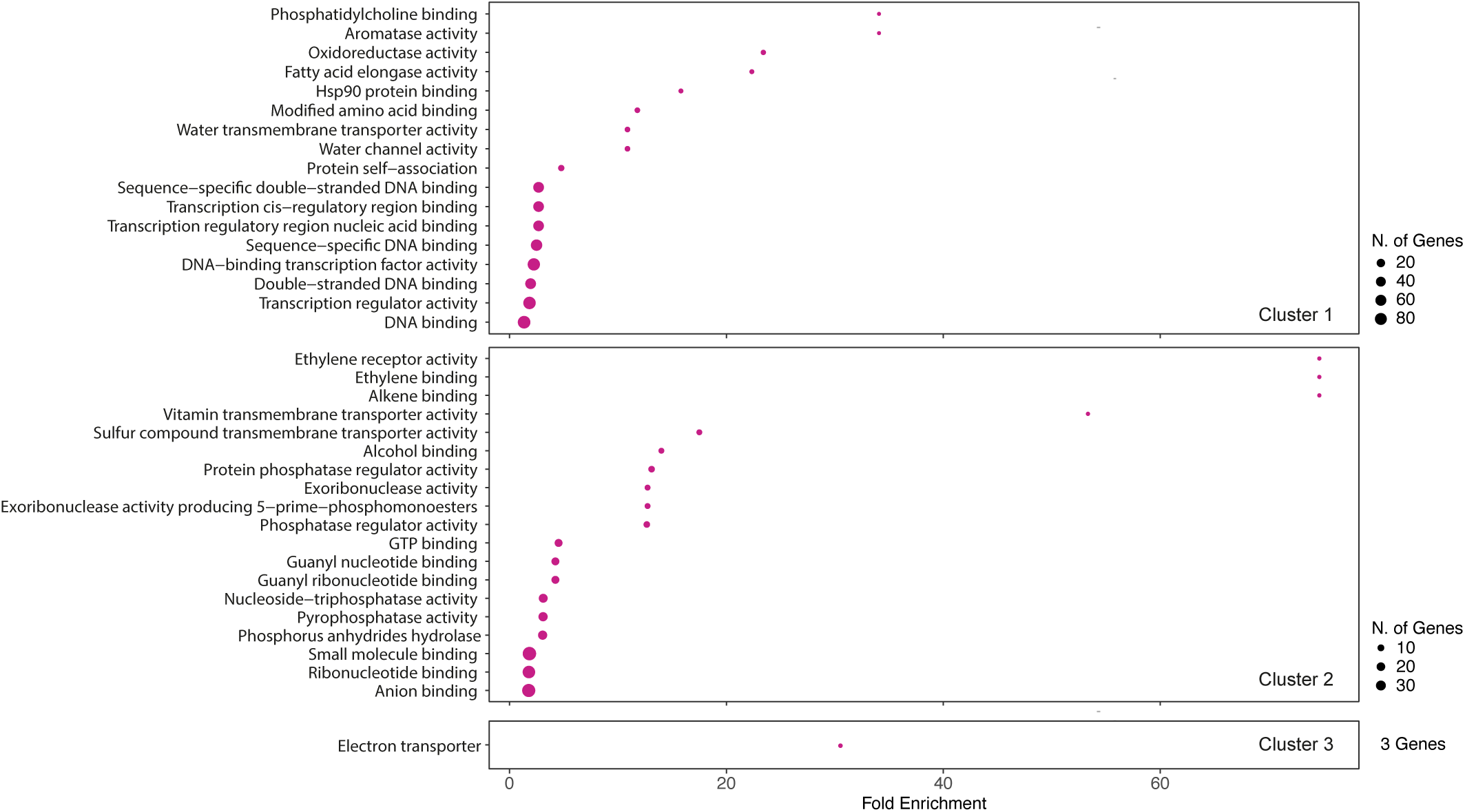
GO enrichment of transcriptional clusters. Gene ontology enrichment analysis of molecular function for the three identified REM associated clusters showing distinct transcriptional dynamics over time (Fig. 1B.).

**Supplementary Figure S4.**
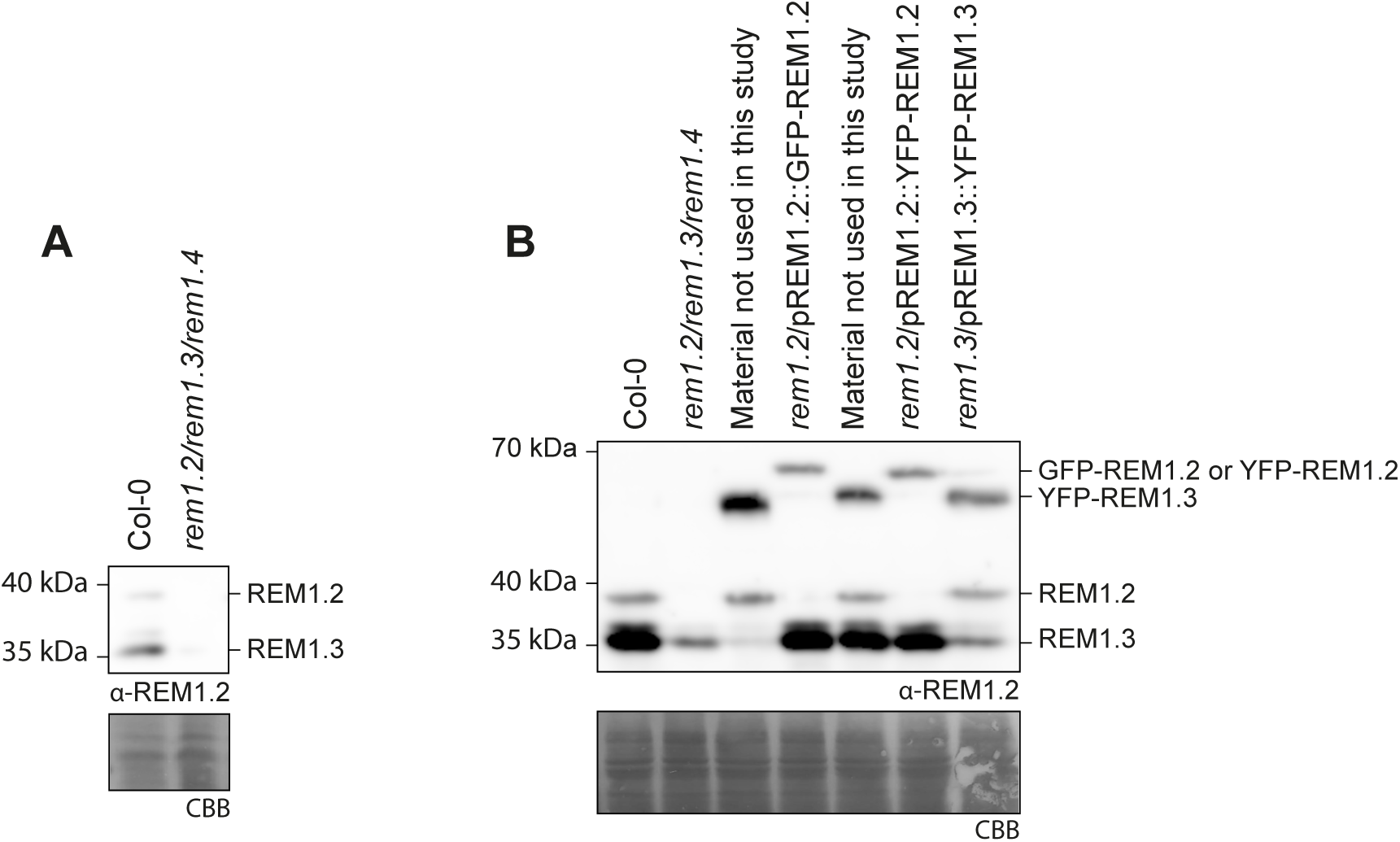
Western blot analysis Arabidopsis transgenic lines. **A-B.** Western blot analysis of REM accumulation in 10 days old seedlings. Western blots were probed with a polyclonal antibody raised against REM1.2, which also recognizes REM1.3. Coomassie brilliant blue (CBB), is presented to show equivalent protein loading.

**Supplementary Figure S5.**
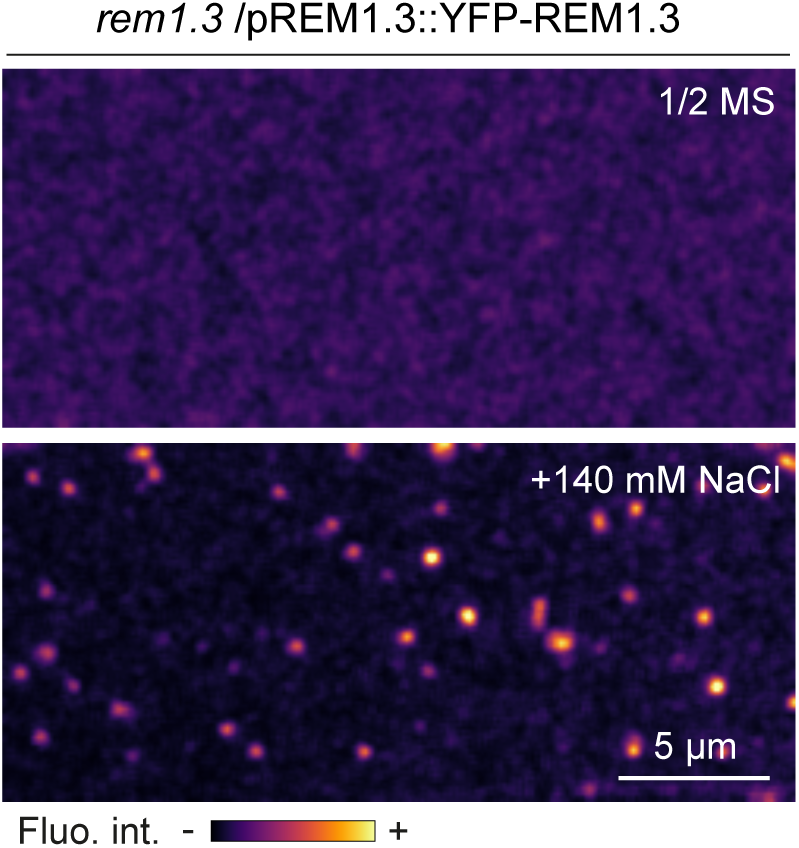
YFP-REM1.3 is reorganized into stable nanodomains upon salt treatment in root epidermal cells. Confocal microscopy pictures of the cell surface of root epidermal cells in presence and in absence of NaCl 140 mM treatment, observed within two minutes of treatment.

**Supplementary Figure S6.**
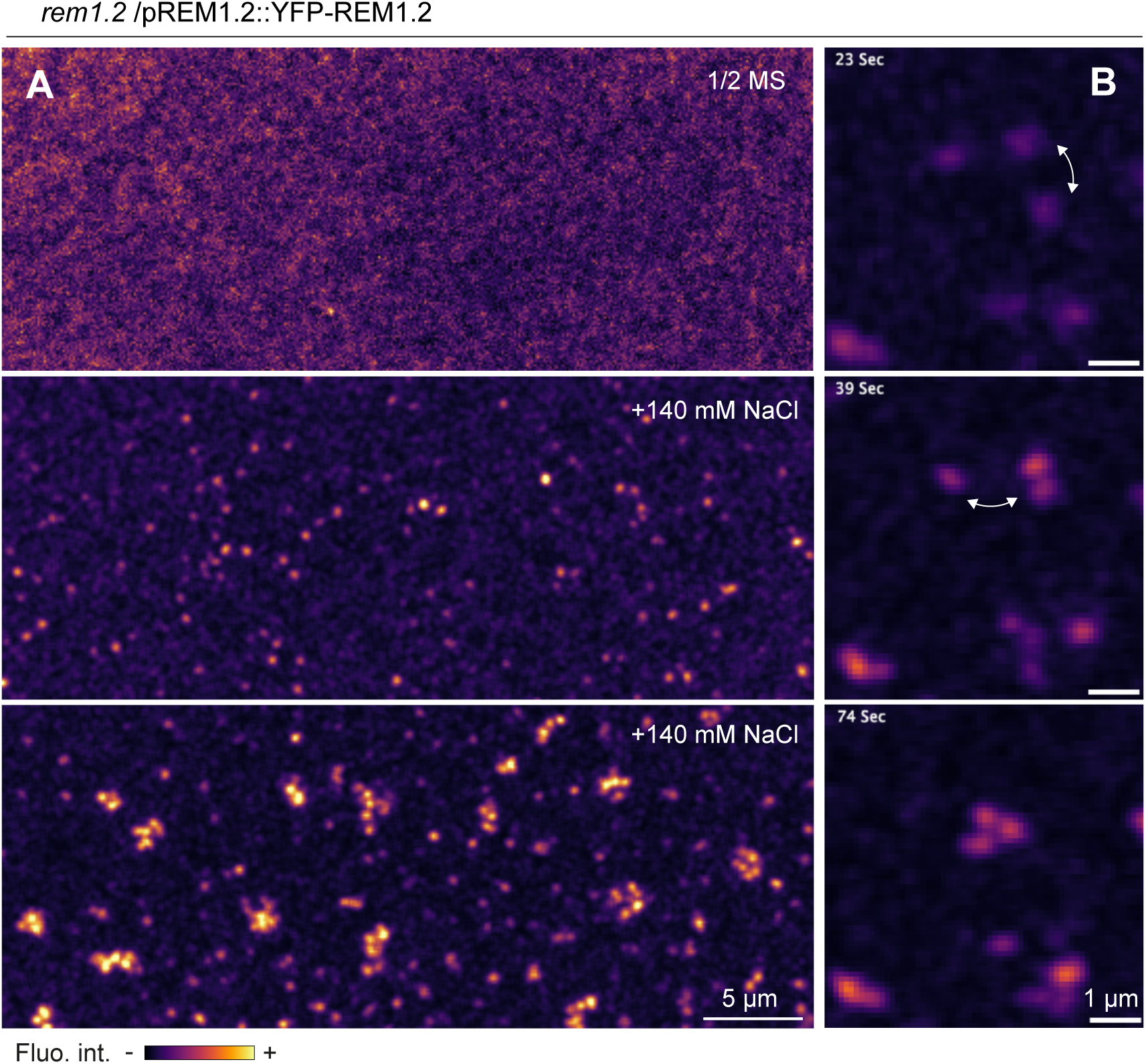
Lateral diffusion of salt-induced YFP-REM1.2 nanodomains in hypocotyl epidermal cells. Confocal microscopy pictures of the cell surface of hypocotyl epidermal cells in presence and in absence of NaCl 140 mM treatment, observed between five and twenty minutes of treatment.

**Supplementary Figure S7.**
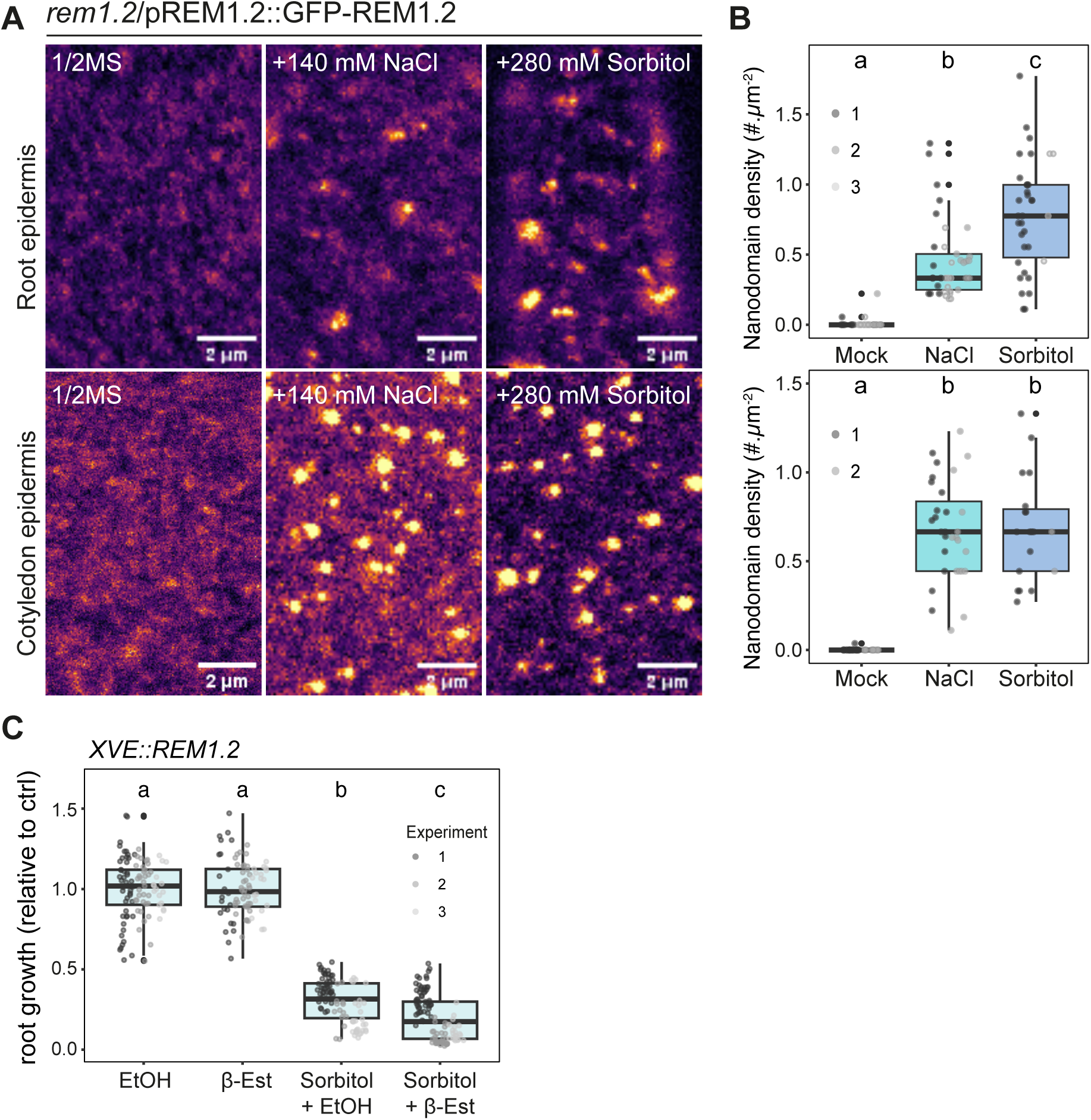
Osmotic stimulation induces GFP-REM1.2 nanodomain organization. **A.** Confocal microscopy pictures of the cell surface of root (top) and cotyledon (bottom) epidermal cells in presence and in absence of NaCl 140 mM and 280mM Sorbitol treatment and imaged between one and five minutes after treatment. **B.** Quantification of GFP-REM1.2 nanodomain density per µm^2^ in root (top) and cotyledon (bottom) epidermal cells. Each dot represents the mean measurement per cell, colors indicate independent experiments, total number of cells (n) in roots mock (n=29), (n=37) for NaCl and (n=35) for Sorbitol and in cotyledons (n=31) for mock, (n=31), for NaCl and (19) for Sorbitol. Between 6 and 15 seedlings per condition were analyzed. Conditions which do not share a letter are significantly different in pairwise Wilcoxon test with Bonferroni p-value adjustment (p< 0,05). **C**. Relative root growth measurements of seedlings transferred to ½ MS containing 0.5 μM β-Estradiol and/or 280 mM Sorbitol and/or corresponding controls for five days. Each dot represents measurements from individual seedlings, colors indicate individual experiments. Measurements are normalized to corresponding mock conditions (without NaCl). Conditions which do not share a letter are significantly different in pairwise Wilcoxon test with Bonferroni p-value adjustment (p< 0,05). Between 79 and 106 roots were measured in total.

**Supplementary Figure S8.**
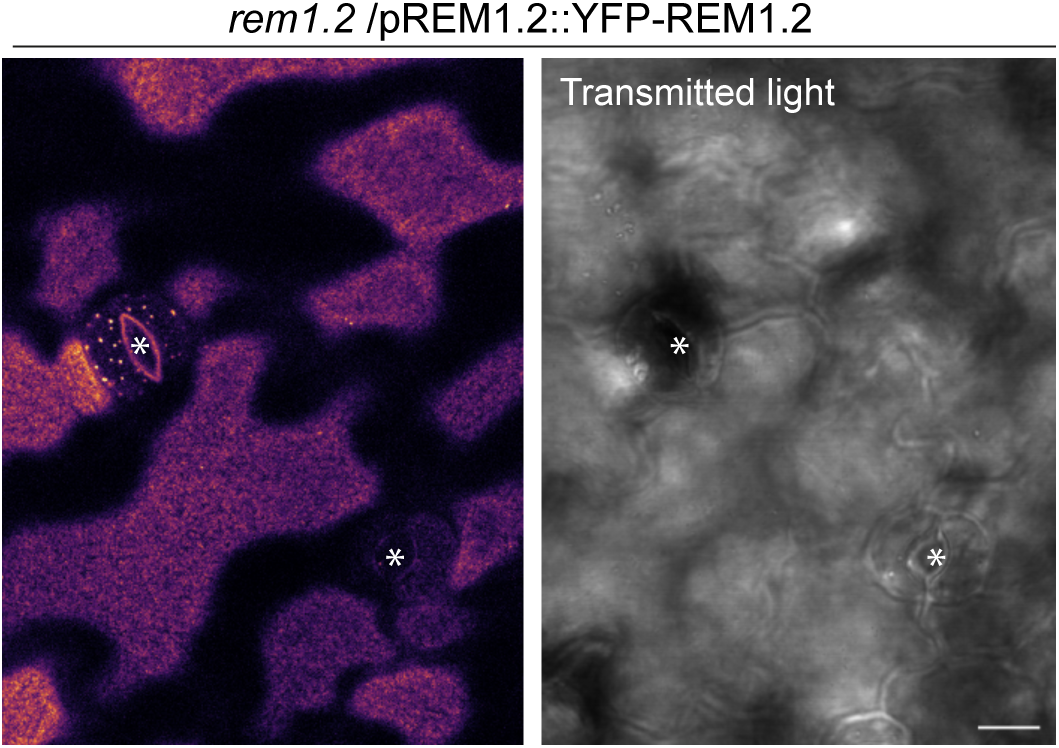
YFP-REM1.2 forms nanodomain in subset of stomata. Confocal microscopy pictures of the cell surface of cotyledon epidermal cells of 5-day-old seedlings with left YFP channel and right Transmitted light. Asterisks indicate Stomata.

**Supplementary Figure S9.**
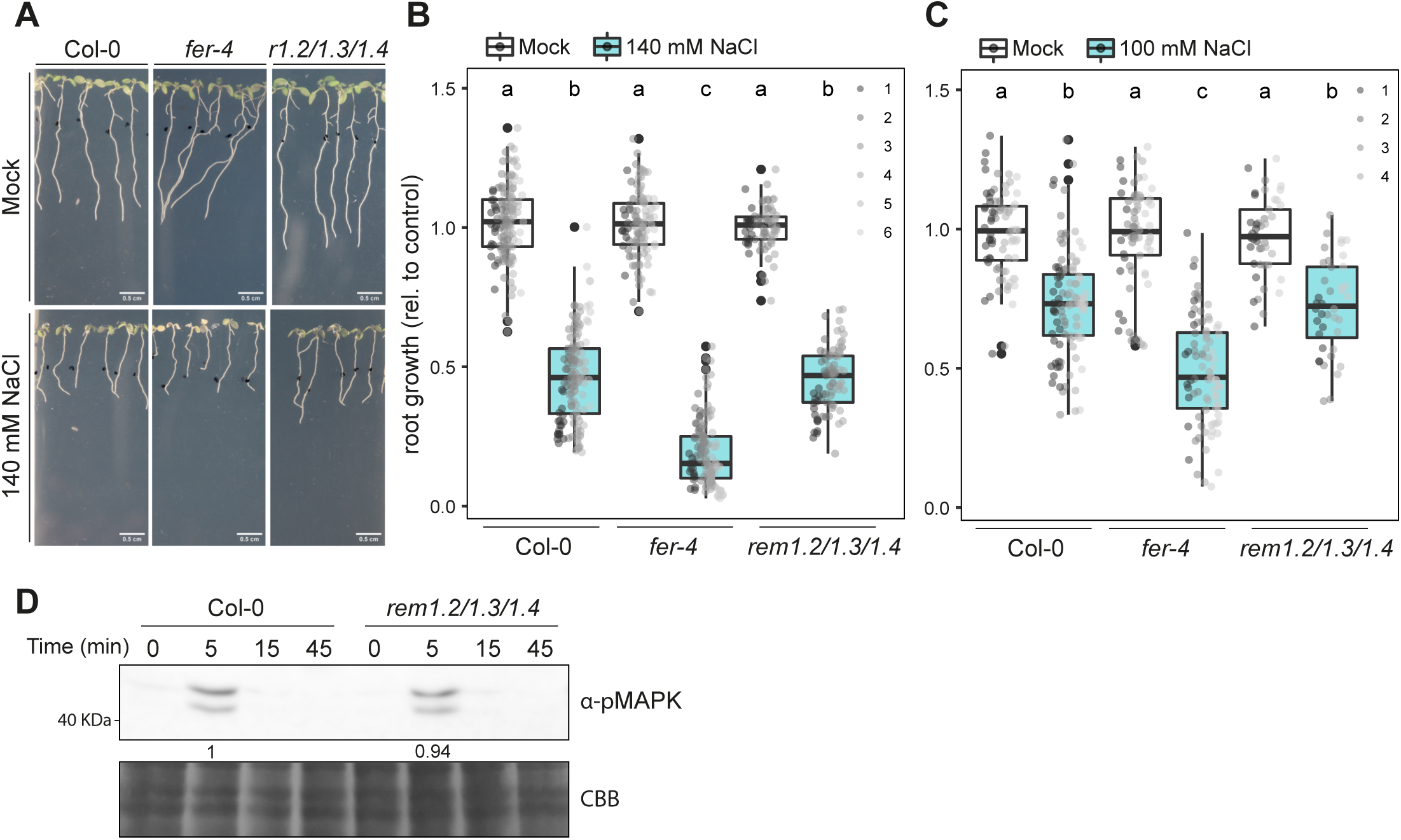
Down regulation of *REM1.2* and *REM1.3* and *REM1.4* does not affect salt signaling and tolerance. **A.** Representative image of plants grown on ½ MS and transferred to ½ MS supplemented with 140 mM NaCl (bottom) or equal amounts of water (top). **B** and **C**. Relative root length measurements of seedlings grown on ½ MS and transferred to ½ MS supplemented with B)140 mM NaCl or C)100 mM NaCl or equal amounts of water (mock). Each dot represents measurements from one individual seedling and dots’ colors indicate individual biological experiments. Measurements are normalized to corresponding mock condition. Conditions which do not share a letter are significantly different in pairwise Wilcoxon test with Bonferroni p-value adjustment (p< 0,05). Total number of seedlings (n) for B) Col-0 mock (n=69), Col-0 NaCl (n=75), fer-4 mock (n=45), *fer-4* NaCl (n=64), *rem1.2/1.3/1.4* mock (n=71), *rem1.2/1.3/1.4* mock (n=81) and for C) Col-0 mock (n=42), Col-0 NaCl (n=57), *fer-4* mock (n=46), *fer-4* NaCl (n=47), *rem1.2/1.3/1.4* mock (n=45), *rem1.2/1.3/1.4* (n=40) **D**. Western blot analysis of salt-induced MAPK phosphorylation in 10-day-old seedlings treated with 140 mM NaCl in time course experiments. Western blots were probed with α-p44/42-ERK revealing phosphorylated MAPKs. Blot stained with Coomassie brilliant blue (CBB), is presented to show equivalent protein loading. The experiment was performed three times with same results.

**Supplementary Figure S10.**
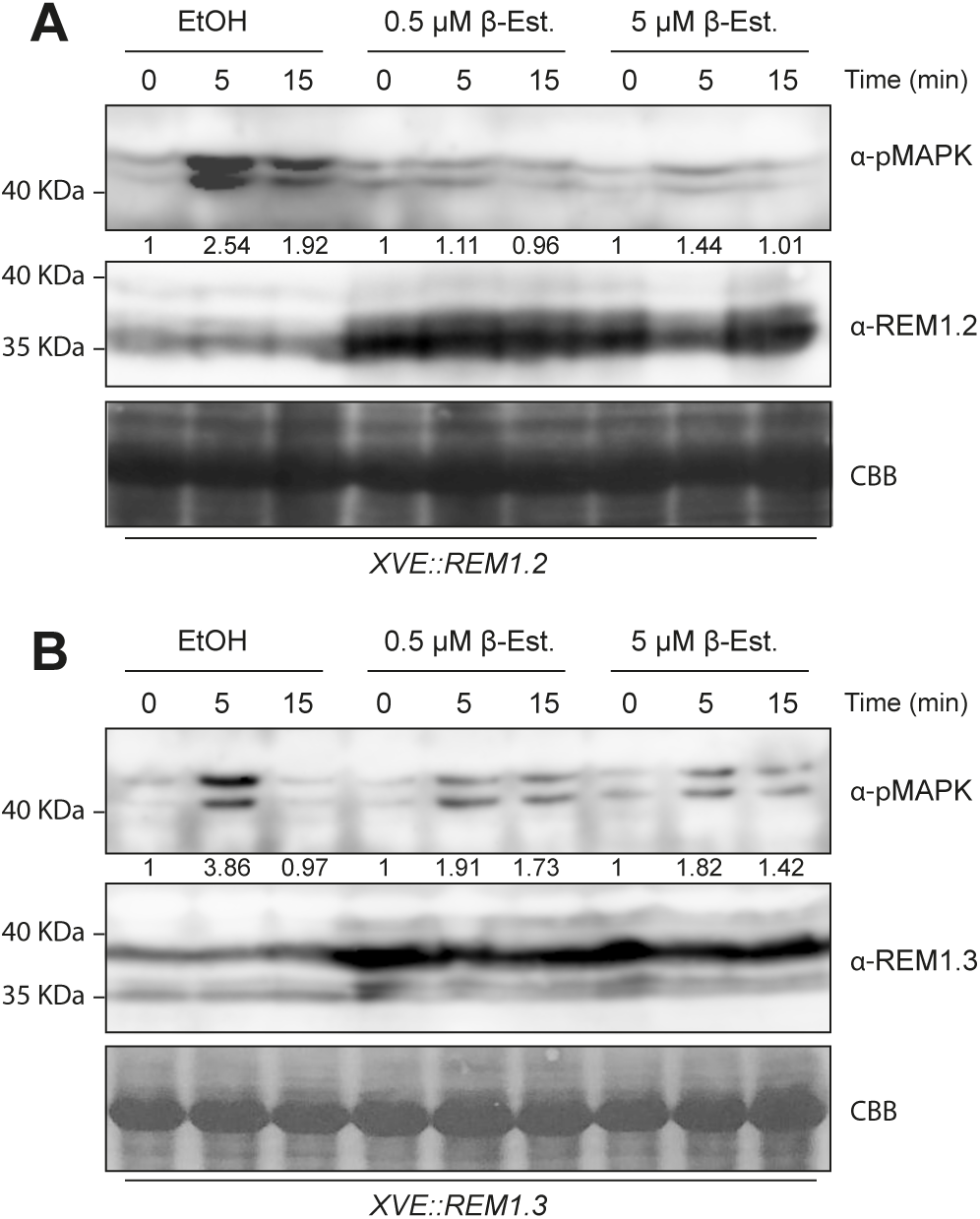
REM1.2 and REM1.3 over-expression inhibits salt-induced MAPK phosphorylation. Western blot analysis of salt-induced MAPK phosphorylation in 10 days old seedlings treated with 140 mM NaCl in time course experiments. Western blots were probed with α-p44/42-ERK revealing phosphorylated MAPKs, α-REM1.2 or α-REM1.3. Blot stained with Coomassie brilliant blue (CBB), is presented to show equivalent protein loading. Similar results were obtained in two independent experiments.

**Supplementary Figure S11.**
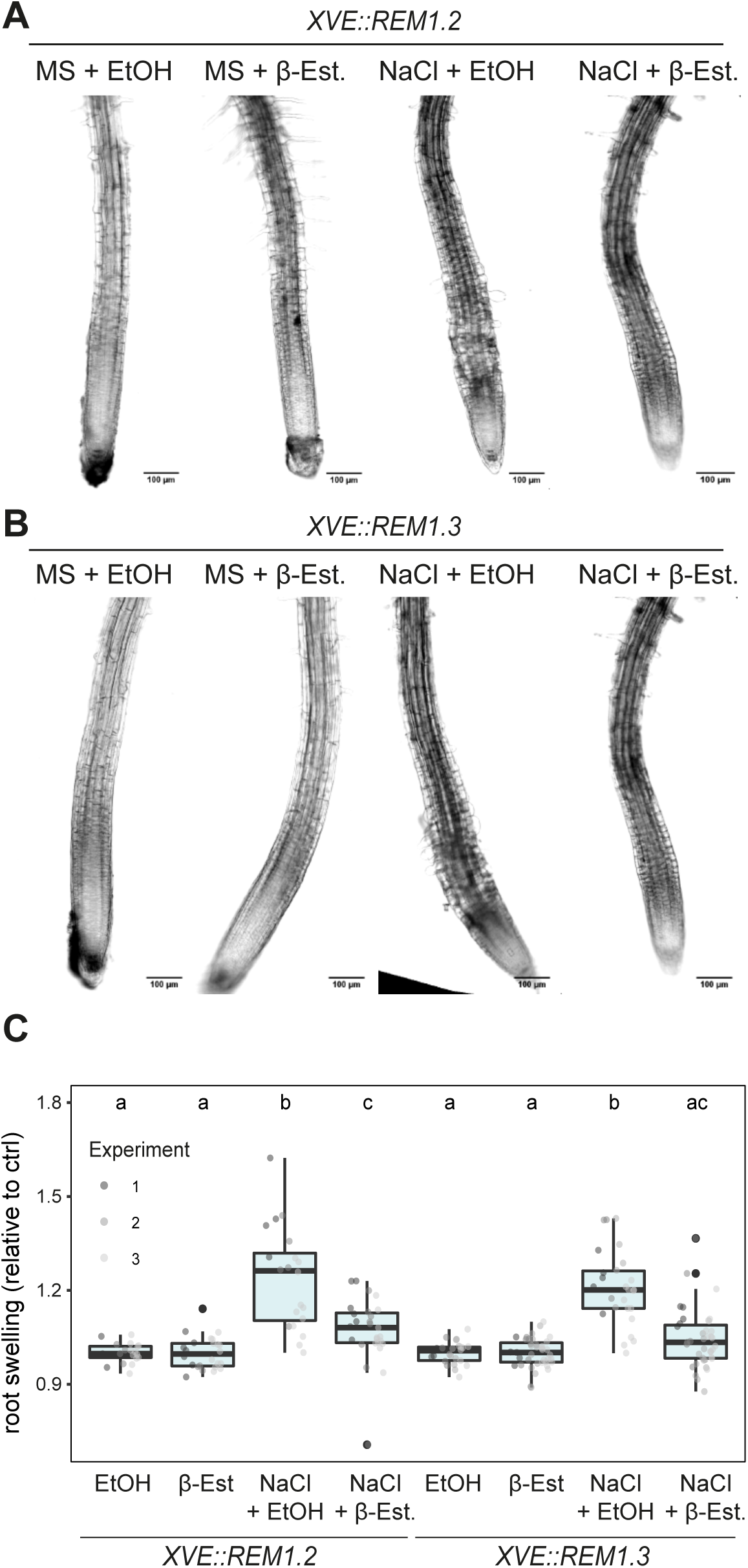
REM1.2 and REM1.3 over-expression inhibits salt-induced root swelling. **A-B**. Representative confocal light transmission images of the root tip of 5 days old seedlings incubated in media 140 mM NaCl in absence and presence of 0.5 μM β-Estradiol. **C**. Quantification of the root swelling of seedlings incubated in liquid ½ MS containing 0.5 μM β-Estradiol and/or 140 mM NaCl and/or corresponding controls for 24h. Each dot represents one measurement from one root, colors indicate independent experiments. Measurements are normalized to corresponding mock conditions (without NaCl). Conditions which do not share a letter are significantly different in pairwise Wilcoxon test with Bonferroni p-value adjustment (p< 0,05). Between 19 and 41 seedlings were analyzed in total.

